# Alpha Cell Theory: Modeling Self-Assembly and Dynamic Processes in Living Systems

**DOI:** 10.1101/2024.11.03.621716

**Authors:** Martin C.H. Gruhlke

## Abstract

Understanding and describing the dynamic processes within living systems is a complex and fundamental challenge. This article presents a novel mathematical model that explores the self-assembly and dynamic behaviors observed in living systems. The model integrates complex chemical reactions, self-organization, and energy transfer processes. Key features of the model include the interaction of various molecular species (A, B, C, D, E, F, G, H, I, and K) and the self-assembly of G-structures within a dynamic environment. Furthermore, the model introduces the concept of Alpha Cell decay when a certain threshold is reached, enhancing the dynamic nature of the system. This research provides insights into the potential underlying mechanisms of life processes and their mathematical representations. It opens the door to a deeper understanding of complex, dynamic, and self-organizing systems in the context of living organisms. The model presented here serves as a valuable tool for further exploration of emergent behaviors and complex dynamics in living systems.

## Introduction

The quest to define life and comprehend its fundamental processes has been a persistent challenge across scientific disciplines. The definition of life, as elusive as it is, has long intrigued biologists, chemists, physicists, and philosophers alike. It remains a question with multiple dimensions, yet its elucidation remains at the heart of our pursuit to understand the essence of living systems [1].

Life, in its myriad forms, exhibits a set of defining characteristics that serve as a starting point for its conceptualization. These characteristics include cellular structure, metabolism, reproduction, growth, adaptation to the environment, and evolution. While these aspects provide a framework, they often prove insufficient in capturing the full scope of life’s complexity, leaving the definition of life an open-ended inquiry [2]. Moreover, life is not solely limited to the biological realm, as the search for extraterrestrial life suggests a broader perspective [3].

In our ongoing exploration of life’s enigma, models play a crucial role in offering insights and simplifications. Theoretical frameworks have emerged, attempting to encapsulate the essence of life in mathematical or conceptual terms. Cellular automata, artificial life simulations, and complexity theory [4,5] are among the tools used to simulate life-like phenomena, shedding light on the self-organizing and emergent behaviors that occur in living systems.

One prominent aspect of life modeling is the concept of dissipative structures. Coined by Ilya Prigogine, dissipative structures refer to systems that are driven far from thermodynamic equilibrium, where they maintain stability and organization by dissipating energy and matter [6]. These structures encompass phenomena such as convection patterns in fluid dynamics, the emergence of life at the molecular level, and the formation of dynamic, self-organizing entities within living systems.

This article presents a novel mathematical model that seeks to explore the intricate interplay of self-assembly, dynamic processes, and energy transfer within living systems. The model, based on complex chemical reactions, aims to encapsulate the essence of life’s emergent properties while integrating the concept of dissipative structures. By examining the formation and decay of self organized G-structures including K molecules (Alpha Cells) within this dynamic environment, this research endeavors to offer a fresh perspective on the mathematical representation of life processes and the mechanisms that underlie self-organization and complex dynamics in living organisms.

In the following sections, we will delve into the specifics of our mathematical model, its theoretical underpinnings, and its potential implications for understanding the complexity of life’s fundamental processes.

### Fundamental Assumptions of the Model

At the core of this model lie a set of foundational assumptions, each of which defines the initial conditions at time t_0_.

We consider a hypothetical scenario taking place in an infinitely large reaction space. This vast environment allows for the virtually unlimited movement and interaction of molecules, enabling an extensive exploration of chemical reactions and self-assembly processes. The system at t_0_ is characterized by the abundant presence of molecular species A, B, E, F, H, and I. These reactants are readily available throughout the infinite reaction space. Their abundant availability ensures that they are not limiting factors in the chemical reactions, promoting continuous and unhindered reactivity.

In contrast, the products C and D, the G-structures, and the catalyst K are absent at t_0_. These components have not yet been formed and introduced into the system, and their emergence will be explored as the model unfolds over time. The environmental conditions, including temperature, pressure, and other relevant factors, are considered constant throughout the entire system at t_0_. These consistent conditions provide a stable backdrop for the chemical reactions and self-assembly processes to occur.

### Chemical Reactions and Characteristics of G and K

In our exploration of dynamic self-assembly within living systems, it is imperative to comprehend the intricate chemical reactions at the heart of the model. This chapter elucidates the reactions themselves and delves into the characteristics of Alpha cells and the catalyst K, both of which play pivotal roles in the model’s evolving narrative.

#### Reaction 1 (Exergonic Reaction): aA + bB -> cC + dD

This exergonic reaction involves the molecular species A and B. Reactants A and B combine in proportions defined by stoichiometric coefficients a and b to yield products C and D, characterized by coefficients c and d. Notably, Reaction 1 is reliant on the presence of the catalyst K, without which it cannot proceed. K, acting as the ignition switch, initiates this exergonic reaction by providing the necessary energy for the transformation of A and B into C and D. This catalytic role of K underscores its significance in the system. Reaction 1 is exergonic, releasing energy as a result of the transformation of A and B into C and D. The energy generated serves as a fundamental source of activation for subsequent processes in the system.

#### Reaction 2 (Endergonic Reaction): eE + fF -> gG

Reaction 2 features the molecular species E and F. Molecules E and F combine in stoichiometric proportions defined by coefficients e and f to form the product G with a coefficient of g. Reaction 2 is endergonic and results in the creation of G-structures. G-structures are dynamic entities with distinctive properties that have a significant impact on the system’s behavior. These structures form barriers within the reaction space, selectively permitting the passage of certain molecules while hindering others. The formation of G-structures, facilitated by Reaction 2, introduces a dynamic element to the system. The properties and behaviors of these structures, such as their size, stability, and effects on molecular diffusion, are central to the model’s dynamics. G-structures are dynamic, bubble-like entities that form as a consequence of physical interactions between G molecules. These structures act as permeable barriers within the reaction space, akin to bubbles in a dynamic fluid medium. They are characterized by the unique property of being selectively permeable, allowing for the passage of specific molecular species, including A, B, C, D, E, F, H, and I, while obstructing the diffusion of K. The continuous integration of new G molecules into the existing structures enhances their growth. G ist at t_0_ not abundant.

#### Reaction 3 (Endergonic Reaction)

Reaction 3 involves molecular species H and I. Molecules H and I react in stoichiometric proportions defined by coefficients h and i to produce the catalyst K, characterized by the coefficient k.

The formation of K in Reaction 3 is crucial, as K plays a multifaceted role in the system. K acts as a catalyst for Reaction 1, enabling the exergonic conversion of A and B into C and D. Additionally, K serves as an essential component for sustaining the system’s energy balance.

Catalyst K is instrumental in initiating Reaction 1, serving as the catalyst that activates the exergonic transformation of A and B into C and D. This catalytic role makes K a central component in the system’s energy generation and subsequent processes. The exergonic nature of Reaction 1, activated by K, releases energy, which is harnessed as an energy source for the system. This energy drives the endergonic processes, including the formation of G-structures in Reaction 2 and the synthesis of additional K in Reaction 3, closing the energy loop. At t_0_ K is also not abundant.

### Local Energy Input at t_1_ to t_2_

At time t_1_, a transient and localized energy input is introduced into the system, setting in motion a cascade of reactions and processes that resonate throughout the reaction space. This input, which extends until t_2_, serves as a crucial driver for the system’s evolution, influencing the rates of various reactions and the self-assembly of G-structures. The energy source fuels the endergonic Reaction 2, responsible for the formation of G-structures.

As the energy infusion commences, the system responds by accelerating Reaction 2, where molecules E and F combine to produce G. This endergonic reaction, stimulated by the energy input, results in the formation of G-structures. These structures, akin to dynamic bubbles, emerge in response to the physical interactions between G molecules.

The distinctive characteristic of this event is that, during the formation of G-structures, the catalyst K, which is a product of Reaction 3, becomes encapsulated within these structures. The encapsulation of K within G-structures has significant implications for the system’s behavior. At this juncture, it is assumed that no K exists outside the boundaries of G-enclosed structures.

The continuous growth and integration of new G molecules into existing structures are fueled by the energy input, enhancing the size of the G-structures. As K becomes confined within these structures, it contributes to the unique dynamics of the system. The localized energy input, the rapid formation of G-structures, and the enclosure of K within them serve as pivotal events that reshape the system’s landscape, heralding a new phase of intricate interactions and emergent phenomena.

This chapter outlines the profound impact of the localized energy input between t_1_ and t_2_ on the formation of G-structures and the encapsulation of K within them. These events mark a critical turning point in the model, setting the stage for the exploration of the complex behaviors and dynamics that arise from this unique interplay.

### Differential Equation for Reaction 1

In this chapter, we delve into the mathematical modeling of Reaction 1, an exergonic reaction that plays a pivotal role in our system. We’ll construct a differential equation that not only takes into account the concentration of K but also considers the diffusion rates of A and B through the G-structures, as well as the diffusion of C and D out of the structures. This reflects the intricate dependencies on reaction constants and diffusion velocities.

The exergonic Reaction 1, which transforms A and B into C and D, can be described using a differential equation that accounts for the concentration of K, the diffusion of reactants, and the diffusion of products. The rate of this reaction, r_1_, is influenced by multiple factors:

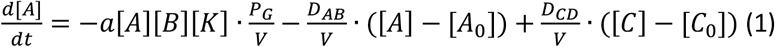

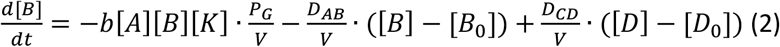

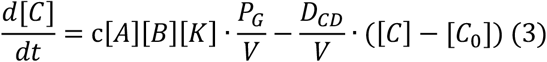

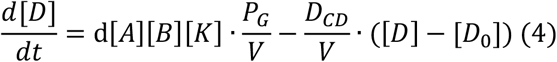

[A] and [B] represent the concentrations of A and B, respectively

[C] and [D] denote the concentrations of C and D.

[K] represents the concentration of [K]

A, b, c and d are the rate constants related to the stochiometric coeffients of the reaction.

P_G_ signifies the permeability of G-structures

V is the volume of the reaction space

D_AB_ represents the diffusion coefficient of A and B through the G structures, assuming that these are the same for A and B.

D_CD_ represents the diffusion coefficient of C and D through the G structures, assuming that these are the same for C and D.

[A_0_], [B_0_], [C_0_], [D_0_] represent the initial concentrations of A, B, C and D, respectively.

The differential equation for Reaction 1 emphasizes the intricate dependencies involved. The reaction rate is determined by the concentration of K ([K]), reaction constants (a, b, c, d), permeability of G-structures (P_G_) and the volume of the reaction space (V). Furthermore, it considers the diffusion of reactants (A and B) through the G-structures, as governed by the diffusion coefficients (D_AB_), as well as the diffusion of products (C and D) out of the structures, as defined by the diffusion coefficients (D_CD_). The concentration changes of these molecules over time are tracked in the differential equations.

This comprehensive approach reflects the rich complexity of Reaction 1, where the transformation of A and B into C and D is intricately linked to reaction constants, diffusion rates, and the role of the G-structures. It sets the stage for understanding how the system’s behavior is shaped by these dynamic interactions, forming the basis for the broader dynamics explored in subsequent chapters.

### Differential Equation for Reaction 2

In this chapter, we will formulate a differential equation for Reaction 2, an endergonic reaction leading to the formation of G-structures. The equation will account for various factors, including the concentration of G, the rates of reactants E and F, and the stability of G-structures.

Reaction 2, characterized by its endergonic nature, results in the formation of G. The rate at which of G production is, *r*_*2*_, is influenced by several factors:

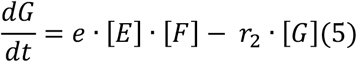

In this equation

[G] represents the concentration G.

[E] and [F] denote the concentration of E and F, respectively,

e is the constant associated with the stoichiometric coefficient of reaction 2

The term -r_2_ [G] describes the rate of G-structure formation, which is contingent on the stability and size of existing G-structures.

The differential equation for reaction 2 highlights the various factors that govern the rate of G-structure formation. These factors include: The concentrations of reactants E and F ([E] and [F]), which are essential components in the formation process. The rate constant e, which determines the reaction kinetics based on the stoichiometric coefficients of reaction 2.

The concentration of G ([G]), which impacts the rate of G-structure formation. Additionally, the equation recognizes that the stability and size of existing G-structures play a role in regulating the rate (r_2_ *[G]). This reflects the dynamic interplay of Reaction 2 within the broader system, where the formation and evolution of G-structures are influenced by the concentrations of reactants, their interaction kinetics, and the characteristics of the G-structures themselves.

The differential equation for Reaction 2 provides a mathematical foundation for understanding the intricate dynamics involved in the endergonic formation of G-structures. It sets the stage for unraveling the system’s behavior in response to localized energy input and the emergence of complex interactions, all of which are explored further in the subsequent chapters.

### Differential equations for reaction 3

In this chapter, we will establish a differential equation for Reaction 3, an endergonic reaction responsible for the synthesis of the essential catalyst K. The equation will consider the concentration of K, the rates of reactants H and I, and the dynamics of K production.

The rate at which K is produced, *r*_*3*_, depends on several factors:

In this equation

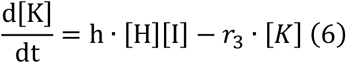

[K] represents the concentration of K

[H] and [I] denote the concentrations of H and I, respectively.

*h* is the rate constant associated with the stoichiometric coefficients of reaction 3.

The term -r_3_ [K] describes the rate of K synthesis, which is contingent on the concentration of H and I.

The differential equation for Reaction 3 underscores the factors that determine the rate of K synthesis. These factors include: The concentrations of reactants H and I ([H] and [I] which are essential components of the synthesis process).

The rate constant *h*, which governs the reaction kinetics based on the stoichiometric coefficients of Reaction 3. The concentration of K ([K]), which influences the rate of K production.

The equation recognizes the dynamic interplay of Reaction 3 within the broader system, where the endergonic synthesis of K is intricately linked to the concentrations of reactants and their interaction kinetics. This dynamic interplay contributes to the evolving behavior of the system.

The differential equation for Reaction 3 serves as a mathematical framework for understanding the complex dynamics involved in the endergonic synthesis of the catalyst K. It lays the foundation for unraveling the system’s response to localized energy input and the emergence of intricate interactions, all of which are explored further in the subsequent chapters.

### Formation of G-structures (Alpha-Cells)

One of the distinctive features of the model ist he dynamic behaviour of G structures. These structures form and evolve as a result of the interactions between G molecules and play a central role in the system’s overall behaviour. Of particular interest ist he behaviour observes, when G structres reach a certain size, after wich they divide into equal parts due to instability of the structure. So the structures grow until they contain a certain number of G molecules, denoted as the treshold value Q. Beyond this threshold, the structures undergo symmetrical division into two identical parts. This treshold has profound implications for the dynamic of the system. **Therefore, alpha cell is defined as a G-structure in which K molecules are enclosed so that a continuous reaction can take place through the influx of reactants and outflow of products**.

Thus, we form a „population” of G structures. The abundance of G structures within the system is not longer solely determined by their growth due to formation of G molecules, but also by their division. As a result, the system becomes more complex and the dynamics of G structures becomes a non-linear pattern.

To describe the growth and division of G structures in the model, one can use a system of differential equations. Hence, we have to introduce two new variabels, Γ and Γ_div_, where Γ represents the number of G strucutres that haven’t reached the treshold number Q of G molecules and Γ_div_ the number of G structures that have divided into two equal parts. As already said, the threshold value for division is denoted as Q.

The rate of increase in the number of G structures that have no reaced the theshold is proportional tot he rate of G moledule formation and the difference between the total numer of G strucutres and the treshold value (Q)

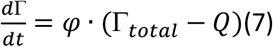

Where:

φ is the rate contant for G-structure formation and

Γ_total_ ist he total number of G-structures.

The rate of increase in the number of G structures that have divided is proportional tot he Rate of G structures reaching the treshold value Q:

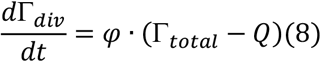

### Example calculation

If we assume that Γ_initla_ = 1 and φ is 0.1 than we can estimate the development of Alphacells as follows:

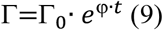

Under these estimations, we find a development of alphacells in an exponential manner, as the graph shows:

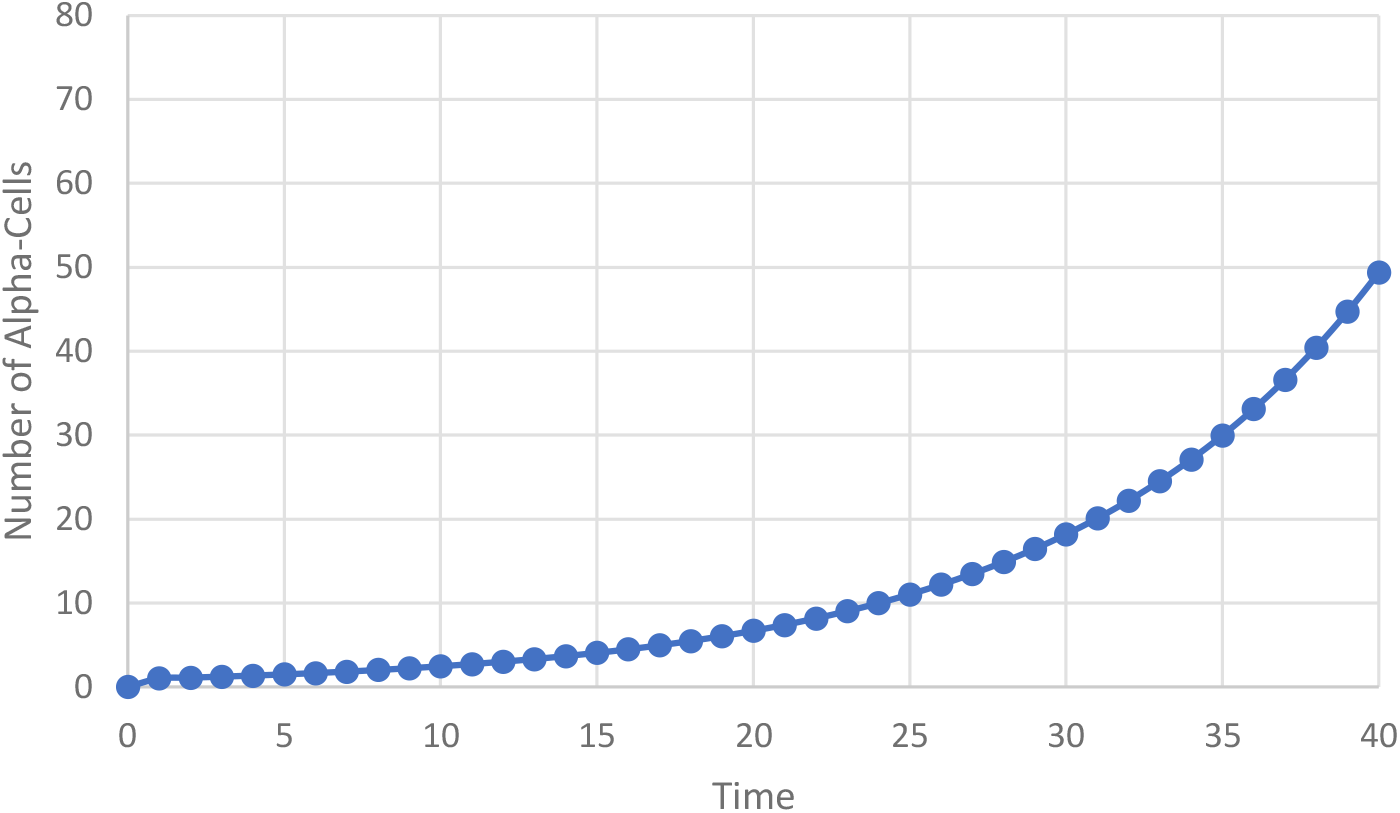

In this exploration of the dynamic model encompassing chemical reactions within an infinitely large compartment, we extend our focus to the emergence and evolution of a population. This population arises from the formation and division of G structures, adding an additional layer of complexity to our chemical ecosystem.

The G structures, formed as a result of boundary molecules in our system, exhibit a fascinating dynamic. As these structures grow, surpassing a predefined threshold, they undergo division, giving rise to new entities. This process mirrors aspects of population growth seen in biological systems. The emergence of a population in our chemical model is intricately linked to the growth and division of these G structures.

The population dynamics in our chemical ecosystem are influenced by the interplay of various factors, including the rate of G structure formation, the division threshold, and the availability of resources represented by molecules A, B, E, F, H, and I. The growth and division of G structures lead to an increase in the overall number of entities within the system, resembling the growth of a population in biological contexts.

The population growth in our model can be mathematically represented by the exponential increase in the number of alpha cells over time. This growth, if sustained, would lead to an ever-expanding population within the chemical compartment. However, the model’s complexity lies in the delicate balance of reactions, ensuring that the emergence of G structures and the subsequent population growth are sustainable within the constraints of available resources.

As our chemical ecosystem evolves, questions of stability and adaptability come to the forefront. The model opens avenues for exploring how variations in reaction rates, threshold values, or the introduction of new elements might impact the stability and evolutionary trajectories of the population. These dynamics bear resemblance to evolutionary processes observed in biological systems, providing an intriguing bridge between chemistry and biology.

The exploration of population dynamics within our chemical model adds a compelling dimension to our understanding of complex, dynamic systems. As we witness the emergence, growth, and potential evolution of a population, the parallels with biological phenomena become evident. This intersection of chemistry and population dynamics invites further inquiry into the fundamental principles governing the emergence of life-like behaviors within chemical systems.

### An attemp in direction of defining life

Defining “life” is a complex and philosophical endeavor, and different perspectives exist depending on the context and field of study. In the context of the dynamic model we’ve explored, incorporating chemical reactions, dissipative structures, and population dynamics, we can offer a nuanced definition:

Life, within the framework of this model, can be conceptualized as an emergent property of dynamic systems exhibiting specific characteristics. These characteristics include the ability to undergo self-sustaining chemical reactions, the emergence of dissipative structures that maintain complexity, and the dynamic evolution of populations within the system.

1. Self-Sustaining Chemical Reactions: Life, in this model, arises from the intricate web of chemical reactions where substances interact and transform, releasing and utilizing energy in a self-sustaining manner.
2. Dissipative Structures: The emergence and maintenance of dissipative structures, exemplified by the G structures, contribute to the organization and complexity within the system. These structures act as boundaries, facilitating specific reactions and processes.
3. Population Dynamics: Life is further characterized by the dynamic evolution of populations, echoing the growth, division, and potential adaptation observed in biological populations. The population dynamics contribute to the overall complexity and adaptability of the system.

In this model, life is not confined to traditional biological entities but extends to dynamic chemical systems capable of sustained reactions, organizational complexity, and population-level behaviors. The definition embraces the idea that life can manifest in diverse forms, not solely dependent on the biochemical composition found in biological organisms.

The model suggests that life exists along a continuum of complexity, where the emergence of dissipative structures and population dynamics marks a transition toward life-like behaviors. This perspective encourages a broader understanding of life as a phenomenon that may manifest in various contexts, not limited to the traditional boundaries of biological organisms.

As we navigate the intersections of chemistry, dissipative structures, and evolving populations, the concept of life becomes a dynamic and evolving phenomenon. Defining life within this framework invites a reevaluation of traditional boundaries and opens new avenues for exploring the fundamental principles that govern the emergence and sustainability of life-like behaviors.

It’s important to note that this definition is specific to the context of the model we’ve discussed, and the broader question of defining life remains a subject of ongoing philosophical, scientific, and interdisciplinary inquiry.

## Conclusion

In this article, we have explored a dynamic model that integrates principles from chemistry, biology, and mathematics to investigate the behavior of chemical reactions within an infinitely large compartment. The model involves reactions between substances A, B, E, F, H, and I, leading to the formation of molecules C, D, G, and K. The presence of a catalytic agent, K, and the formation of a boundary molecule, G, contribute to the complexity of the system.

The system’s behavior is characterized by the exergonic reaction of A and B, the endergonic reactions of E and F, and H and I, and the catalysis provided by K. The emergence of G structures, forming a permeable barrier, adds an additional layer of intricacy to the system. These structures exhibit growth until reaching a specified threshold, at which point they divide into two equal parts.

Our mathematical analysis has shown that the growth of G structures, when not impeded by division, follows an exponential pattern. However, the concentrations of molecules C, D, and K are influenced by multiple factors, including the stoichiometry of reactions and the availability of reactants.

This model offers insights into dissipative structures in chemical systems, highlighting the interplay between exergonic and endergonic reactions, catalysis, and the emergence of boundary structures. Further refinement of the model, incorporating specific rate equations and constants, will enable a more accurate representation of real-world chemical systems.

As we delve deeper into understanding the dynamics of chemical reactions within vast compartments, the implications for defining life become more profound. The delicate balance of reactions, energy transfer, and the emergence of dissipative structures raises intriguing questions about the fundamental nature of life and the conditions that foster its emergence.

In conclusion, this model provides a foundation for future research at the intersection of chemistry, biology, mathematics and philosphy. By refining and expanding upon these concepts, we may gain deeper insights into the principles governing the emergence and dynamics of life in complex chemical systems.

